# Role of β-arrestin-2 in short- and long-term opioid tolerance in the dorsal root ganglia

**DOI:** 10.1101/2020.11.15.383620

**Authors:** Karan H. Muchhala, Joanna C. Jacob, William L. Dewey, Hamid I. Akbarali

## Abstract

β-arrestin-2 has been implicated in the mechanism of opioid-induced antinociceptive tolerance. G-protein-biased agonists with reduced β-arrestin-2 activation are being investigated as safer alternatives to clinically-used opioids. Opioid-induced analgesic tolerance is classically considered as centrally-mediated, but recent reports implicate nociceptive dorsal root ganglia (DRG) neurons as critical mediators in this process. Here, we investigated the role of β-arrestin-2 in the mechanism of opioid tolerance in DRG nociceptive neurons using β-arrestin-2 knockout mice and the G-protein-biased μ-opioid receptor agonist, TRV130. Whole-cell current-clamp electrophysiology experiments revealed that 15-18-hour overnight exposure to 10 μM morphine *in vitro* induced acute tolerance in β-arrestin-2 wild-type but not knockout DRG neurons. Furthermore, in wild-type DRG neurons circumventing β-arrestin-2 activation by overnight treatment with 200 nM TRV130 attenuated tolerance. Similarly, in β-arrestin-2 knockout male mice acute antinociceptive tolerance induced by 100 mg/kg morphine s.c. was prevented in the warm-water tail-withdrawal assay. Treatment with 30 mg/kg TRV130 s.c. also inhibited antinociceptive tolerance in wild-type mice. Alternately, in β-arrestin-2 knockout DRG neurons tolerance induced by 7-day *in vivo* exposure to 50 mg morphine pellet was conserved. Likewise, β-arrestin-2 deletion did not mitigate *in vivo* antinociceptive tolerance induced by 7-day exposure to 25 mg or 50 mg morphine pellet in both female or male mice, respectively. Consequently, these results indicated that β-arrestin-2 mediates acute but not chronic opioid tolerance in DRG neurons and to antinociception. This suggests that opioid-induced antinociceptive tolerance may develop even in the absence of β-arrestin-2 activation, and thus significantly affect the clinical utility of biased agonists.

## 1. Introduction

μ-opioid receptors (MORs) are known to engage scaffolding proteins called β-arrestins, which classically function to desensitize activated GPCRs through stearic inhibition, but can also activate molecular mechanisms independent of G-protein signaling (Lefkowitz, 1998; Lohse et al., 1990; Williams et al., 2013). Previously, studies have shown that genetic deletion or down-regulation of β-arrestin-2, or use of G-protein-biased agonists enhances antinociception and reduces antinociceptive tolerance in rodents, thus implicating β-arrestin-2 in the mechanism of opioid-induced antinociceptive tolerance (Bohn et al., 2000, 1999; DeWire et al., 2013; Grim et al., 2020; Manglik et al., 2016; Wang et al., 2016; Yang et al., 2011).

Primary afferent neurons of the dorsal root ganglia (DRG), specifically the thinly myelinated Aδ- and unmyelinated C-fiber neurons, are first-order components of the ascending pain pathway. These neurons generate and transduce chemical, mechanical and thermal noxious stimuli as action potentials from the periphery to second-order neurons in the spinal cord, which synapse with neurons in the pain center in the brain and with motor neurons of the spinal reflex arc (Stein and Machelska, 2011). Thus, nociceptive DRG neurons generate signals that eventually get processed in the CNS as “pain”.

While the prevailing dogma is that opioid-induced analgesia and tolerance are centrally mediated, recent studies have highlighted the critical role of nociceptive DRG neurons in the expression of antinociception and induction of opioid tolerance. Selective deletion of MORs from Nav1.8-containing DRG neurons diminished the ability of morphine to mitigate inflammatory pain (Weibel et al., 2013), whereas elimination of MORs from the entire population of DRG neurons abolished the acute spinal and supraspinal antinociceptive effects of opioids (Sun et al., 2020, 2019). Ablation of TRPV1-expressing DRG neurons or conditional knockout of MORs on TRPV1-expressing DRG neurons reversed antinociceptive tolerance (Chen et al., 2007; Corder et al., 2017), suggesting that MORs on primary afferent DRG neurons could be important targets for the prevention of antinociceptive tolerance. Therefore, it is important to delineate the molecular mechanisms underlying cellular tolerance in DRG nociceptive neurons, specifically the role of β-arrestin-2.

In the present study, we investigated the role of β-arrestin-2 in the mechanism of opioid tolerance in small-diameter DRG nociceptive neurons using β-arrestin-2 knockout mice and the G-protein-biased MOR agonist, TRV130 (DeWire et al., 2013). We demonstrate that acute or “short-term” tolerance—manifested as a result of several hours (overnight) of opioid exposure (Williams et al., 2013)—in nociceptive DRG neurons is mediated by β-arrestin-2, whereas chronic or “long-term” tolerance—developing due to seven days of opioid exposure (Williams et al., 2013)— is independent of β-arrestin-2. Acute antinociceptive tolerance to *in vivo* exposure to TRV130 or morphine in mice for one day is also attenuated in the absence of β-arrestin-2 activation. However, chronic antinociceptive tolerance to morphine in either male or female mice develops independent of the β-arrestin-2 pathway. In conclusion, the findings presented in this study indicate that β-arrestin-2 is a critical mediator of acute tolerance but does not underlie chronic tolerance in DRG nociceptive neurons.

## 2. Materials and Methods

### 2.1. Drugs and Chemicals

Morphine sulfate pentahydrate and implantable morphine pellets (25 mg) were obtained from the National Institutes of Health National Institute on Drug Abuse (Bethesda, MD). (+) TRV130 was obtained from Dr. Bruce Blough (Research Triangle Institute, NC). Pyrogen-free isotonic saline was purchased from Hospira (Lake Forest, IL). Dulbecco’s modified Eagle’s medium (DMEM)/F12, Neurobasal-A medium, Ca^2+^ and Mg^2+^ -free Hank’s balanced salt solution (HBSS), 50x B-27 supplement and L-glutamine were purchased from Gibco, Thermo Fisher Scientific (Waltham, MA). Penicillin/streptomycin/amphotericin B antibiotic-antimycotic solution and laminin were purchased from Corning (Corning, NY). Papain, glial cell line-derived neurotrophic factor (GDNF) and fetal bovine serum (FBS) were purchased from Worthington Biochemical Corporation (Lakewood, NJ), Neuromics (Edina, MN) and Quality Biological, (Gaithersburg, MD), respectively. Glass cover slips were purchased from ThermoFisher Scientific (Waltham, MA). Twenty-four-well culture dishes and 35 × 10 mm petri dishes were purchased from CELLTREAT (Pepperell, MA). Collagenase from *Clostridium histolyticum*, poly-D-lysine, CaCl_2_, MgCl_2_, NaCl, KCl, HEPES, EGTA, NaH_2_PO_4_, glucose, Na_2_ATP, NaGTP, L-aspartic acid (K salt), KOH, NaOH, MgSO_4_ and NaHCO_3_ were purchased from MilliporeSigma (Burlington, MA).

### 2.2. Animals

All animal care and experimental procedures were conducted in accordance with procedures reviewed and approved by the Institutional Animal Care and Use Committee at Virginia Commonwealth University in compliance with the US National Research Council’s Guide for the Care and Use of Laboratory Animals, the US Public Health Service’s Policy on Humane Care and Use of Laboratory Animals, and Guide for the Care and Use of Laboratory Animals.

Male and female β-arrestin-2 (βarr2) knockout (KO) or their wild-type (WT) littermates were separately group-housed by genotype with up to five animals per IVC cage in animal-care quarters maintained at 22°C ± 2°C on a 12-hour light/dark cycle. Mice used in experimental procedures were at least 7 weeks of age. Male β-arrestin-2 mice weighed 20-25 g, and female β-arrestin-2 mice weighed 19-23 g. Mice were acclimated in the vivarium for at least one week prior to experimentation. Breeding pairs for the β-arrestin-2 mice were initially obtained from Dr. Lefkowitz (Duke University, Durham, NC) and housed within the transgenic facility at Virginia Commonwealth University. The genetic background of the β-arrestin-2 WT and KO mice used in our experimental procedures was 81% C57B6J:19% C57B6N. Mice had access to food and water *ad libitum* unless specified otherwise.

### 2.3. *In vivo* morphine treatment

Subcutaneously implanted continuous-release morphine pellets were used to model chronic in vivo exposure as pellets maintain a high plasma level of morphine over a longer period of time compared to intermittent injections or osmotic pumps and the tolerance induced is more robust (Dighe et al., 2009; McLane et al., 2017). Where indicated, one or two 25 mg morphine pellets were implanted subcutaneously in the dorsum of female or male β-arrestin-2 mice for 7 days, respectively. This dose was based on a previous study, where we demonstrated that a 7-day exposure to one 25 mg and two 25 mg morphine pellets in female and male β-arrestin-2 mice, respectively, substantially right-shifted the morphine dose response curve compared to wild-type mice in the warm-water tail-withdrawal assay, indicating antinociceptive tolerance (Muchhala et al., 2020). The dose of two 25 mg morphine pellets is henceforth denoted as “50 mg” in the present study. In all experiments, control mice were implanted with one or two placebo pellets. In order to implant the pellet, mice were first anesthetized with 2.5% isoflurane before shaving the hair from the base of the neck. Skin was disinfected with 10% povidone iodine (General Medical Corp, Walnut, CA) and alcohol. A 1 cm horizontal incision was made at the base of the neck and one or two pellets were inserted in the subcutaneous space. The surgical site was closed with Clay Adams Brand, MikRon AutoClip 9-mm wound clips (Becton Dickinson, Franklin Lakes, NJ) and cleansed with 10% povidone iodine. Use of aseptic surgical techniques minimized any potential contamination of the pellet, incision and subcutaneous space. Mice were allowed to recover in their home cages where they remained throughout the experiment.

A repeated injection schedule as described previously by Bohn et al. (Bohn et al., 2002) was also used to induce acute antinociceptive tolerance. Here, mice were injected subcutaneously with a high dose of morphine (100 mg/kg s.c.) or TRV130 (30 mg/kg s.c.) on Day 1. Control mice received saline. On the next day, antinociceptive tolerance was assessed using the warm-water tail-withdrawal assay (as described below) by injecting mice with either 10 mg/kg morphine s.c. or 3 mg/kg TRV130 s.c. The doses of TRV130 used in this experiment were adjusted to their morphine equivalents.

### 2.4. Evaluating thermal nociception

Thermal nociception was examined using the warm-water tail-withdrawal test, which represents the sensory aspects of spinally-mediated acute pain and has been classically used to test the efficacy of opioid analgesics (Mogil, 2009). In the warm-water tail-withdrawal assay, mice were gently secured in a cloth and the distal 1/3^rd^ of the tail was immersed in a water bath warmed to 56°C ± 0.1°C. The latency to withdraw the tail from the water was recorded. A maximum cut-off of 10 seconds was set to prevent damage to the tail. Only naïve mice with control latency between 2 and 4 seconds were used in experiments. Tolerance was assessed by challenging mice with an acute subcutaneous injection of morphine or TRV130. Challenge latency was compared against baseline latency. Where indicated, antinociception was quantified as %MPE, which was calculated as follows: %MPE= [(Challenge latency-baseline latency)/ (10-baseline latency)] × 100, adapted from Harris and Pierson, 1964 (Harris and Pierson, 1964).

### 2.5. Behavioral testing

All testing was conducted in a temperature and light-controlled room in the light phase of the 12-hour light/dark cycle. Mice were acclimated to the testing room for at least 15-18 hours before commencing experiments to mitigate stress to the animals and eliminate confound from potential stress-induced effects on antinociception (Sorge et al., 2014). All animals were randomly divided into control and treatment groups. Mice were excluded from experiments if they exhibited wounds from aggressive interactions with cage mates, since injury-induced activation of the endogenous opioid system could confound nociceptive assays (Corder et al., 2013).

### 2.6. Isolation and primary culture of dorsal root ganglia neurons

Dorsal root ganglia (DRG) cells were prepared from adult mice as previously described (Ross et al., 2012). Mice were sacrificed via CO_2_ inhalation and L5 – S1 DRGs were immediately harvested using a dissecting microscope. DRGs were placed in a 35 mm dish containing Hank’s balanced salt solution (HBSS) and papain (15 U/ml) was added prior to incubation at 37°C for 18 minutes. After the initial incubation step, ganglia were transferred to a new 35 mm dish containing HBSS and 1.5 mg/ml collagenase from *Clostridium histolyticum* and incubated for 1 hour at 37°C. DRGs were then transferred to a sterile 15 ml conical tube containing ice-cold (4°C) DMEM/F12 supplemented with 10% FBS and dissociated by trituration before being centrifuged at 350 × g for 5 minutes. The supernatant was decanted and the pellet was resuspended in neurobasal A media that contained 1% FBS, 1x B-27 supplement, 10 ng/ml GDNF, 2 mM L-glutamine, and 100 U/ml penicillin/streptomycin/amphotericin B (complete neuron media). Cells were then plated on glass cover slips (1/well) coated with laminin and poly-D-lysine. Twenty-four-well plates containing isolated DRGs were then incubated overnight at 37°C in a humidified 5% CO_2_/air stabilized incubator. Whole-cell patch clamp electrophysiology experiments were conducted 15-18 hours later on the following day. Where indicated, neurons were exposed to 10 μM morphine sulfate pentahydrate or 200 nM TRV130 in complete neuron media for 15-18 hours (overnight) prior to whole-cell patch clamp experiments to mimic prolonged opioid exposure.

Cells were also isolated from male β-arrestin-2 WT or KO mice subcutaneously implanted with 50 mg morphine pellet for 7 using the procedure described above. Isolated cells were incubated as described above in Neurobasal-A media supplemented with growth factors and antibiotic-antimycotic solution for 15-18 hours.

### 2.7. Whole-cell patch clamp electrophysiology

Micropipettes for patch clamp experiments (2-4 MΩ) were made from pulled (Model P-97 Flaming-Brown Micropipette Puller, Sutter Instruments, Novato, CA) and fire-polished 1.5/0.84 o.d./i.d. (mm) borosilicate glass capillaries (World Precision Instruments, Sarasota, FL). The internal physiological solution was composed of 100 mM L-aspartic acid (K salt), 30 mM KCl, 4.5 mM Na_2_ATP, 1 mM MgCl_2_, 10 mM HEPES, 0.1 mM EGTA and 0.5 mM NaGTP and pH adjusted to 7.2 with 3 M KOH. Coverslips with adherent cells were transferred to a microscope stage plate continuously superfused with external physiological salt solution composed of 135 mM NaCl, 5.4 mM KCl, 0.33 mM NaH_2_PO_4_, 5 mM HEPES, 5 mM glucose, 2 mM CaCl_2_ and 1 mM MgCl_2_, and adjusted to a pH of 7.4 with 1 M NaOH. Whole-cell current-clamp recordings were made in a room temperature environment using HEKA EPC 10 (HEKA, Bellmore, NY) or Axopatch 200B (Molecular Devices, Sunnyvale, CA) amplifiers at a 10 kHz sampling frequency and 2.9 kHz low-pass Bessel filtering. Pulse generation and data acquisition were achieved with PatchMaster v2×60 (HEKA) or Clampex and Clampfit 10.2 software (Molecular Devices). The current-clamp protocol consisted of 10 pA steps, beginning at −30 pA to assess both active and passive cell properties. All current clamp recordings were performed two-five minutes after achieving whole-cell mode to allow dialysis of internal solution. Pre- “baseline” recordings were taken to ensure cell stability before beginning drug perfusion. Once neurons were deemed stable, the external solution source was exchanged for the external solution containing 3 μM morphine or 50 nM TRV130. A recording was taken immediately following this exchange, and was labeled “T_0_” or time zero, which served as the “baseline” recording in all studies. A recording was then taken for up to 16 minutes to capture neuronal responses to morphine or TRV130. Electrical properties such as threshold potential (V_thresh_), rheobase, resting membrane potential (V_rest_) and input resistance (R_input_) were extrapolated from each recording, and the difference between baseline and following drug exposure was calculated for each cell. Action potential (AP) derivatives were determined using the differential function in the PatchMaster or Clampfit software, where the derivative of the voltage with respect to time (dV/dt) was calculated in order to estimate threshold potential. Threshold potential was defined as the voltage at which dV/dt significantly deviated from zero during the course of the action potential uprise. It was used as the primary measure of neuronal excitability in our experiments. Each coverslip was discarded following drug exposure and the same process was repeated on a freshly-mounted coverslip. Cellular tolerance was assessed identically in cells either incubated with 200 nM TRV130 or 10 μM morphine overnight, or isolated from male mice implanted with two 25 mg morphine pellets for 7 days. Values reported were not corrected for junction potentials (~12 mV).

Experiments were performed only on cells with healthy morphology and stable patch. In DRG primary isolations, small-diameter neurons (< 30 pF capacitance) correspond to nociceptive Aδ fiber and C-type neurons (Abraira and Ginty, 2013; Barabas et al., 2014), therefore we preferentially selected these cells for use in our experiments. Due to the presence of multiple subtypes of small-diameter DRG neurons in our primary cultures, measures from individual neurons were considered as independent values and not replicates for data analysis. In all electrophysiology experiments ‘n’ represents the number of neurons per group and ‘N’ denotes number of animals per group.

### 2.8. Data and statistical analysis

Data analysis was performed in GraphPad Prism 8.0 (GraphPad Software, Inc., La Jolla, CA). Data are expressed as mean ± S.E.M. Depending on the experimental design data were analyzed using 2-tailed unpaired Student’s t-test, 2-way ANOVA or 2-way repeated measures ANOVA. Select data were also analyzed by multiple 2-tailed paired t-tests with two-stage step-up method of Benjamini, Krieger and Yekutieli, and a false discovery rate of 5%. Where specified in electrophysiology experiments, 3-way repeated-measures ANOVA was used to evaluate the main effect of three independent variables on action potential threshold, rheobase, resting membrane potential or input resistance. Bonferroni’s multiple comparisons post-hoc test was conducted only if the F value in the ANOVA table was significant (Curtis et al., 2018). An alpha level of 0.05 was pre-determined. Thus, data were statistically significant when P <0.05. The specific statistical tests used for data analysis are indicated in the text or figure legends. Sample sizes were based on our previous studies with similar experimental protocols.

## 3. Results

### 3.1. β-arrestin-2 knockout prevents the development of short-term tolerance in DRG neurons

We investigated whether opioid tolerance in DRG nociceptors following overnight (short-term) exposure is mediated by β-arrestin-2. For this purpose, we harvested DRG neurons from male β-arrestin-2 WT or KO mice and incubated them in media containing 10 μM morphine for 15-18 hours (overnight). Cellular tolerance was assessed in only small-diameter DRG neurons (membrane capacitance < 30 pF) by challenging them with 3 μM morphine (Fig. 1). We used threshold potential, which is the membrane potential at which action potential is elicited, as the primary metric of neuronal excitability. An increase in threshold potential from baseline represented reduced neuronal excitability, whereas a cell was described as “tolerant” if the acute challenge had no effect on threshold potential. In untreated DRG neurons from both β-arrestin-2 WT and KO mice, acute 3 μM morphine significantly increased action potential threshold from baseline (WT: −16.5 ± 2.1 mV to −11.4 ± 2.0 mV, P<0.001; KO: −18.4 ± 2.8 mV to −14.1 ± 2.5 mV, P<0.001; Figs. 1A, 1B and 1C; Table S1). In contrast, no significant shift in action potential threshold from baseline was observed when WT neurons previously exposed to 10 μM morphine overnight were challenged with 3 μM morphine (−18.2 ± 1.5 mV to −17.9 ± 1.8 mV, P>0.05; Figs. 1A and 1C; Table S1). The shift in threshold potential from baseline was 0.2 ± 0.4 mV, which is significantly decreased compared to the shift of 5.0 ± 1.1 mV in naïve βarr2 WT neurons (P<0.01; Fig. 1D). These data together signify the development of cellular tolerance in overnight morphine-treated WT neurons. However, unlike WT neurons, βarr2 KO DRG neurons exhibited reduced threshold potentials in response to the acute 3 μM morphine challenge despite being exposed to 10 μM morphine overnight (−19.9 ± 3.0 mV to −15.5 ± 3.9 mV, P<0.001; Figs. 1B and 1C; Table S1). The shift in threshold potential of 4.4 ± 1.1 mV in overnight morphine-treated βarr2 KO neurons was not significantly different from the threshold potential change of 4.3 ± 0.8 mV in naïve βarr2 KO neurons (P>0.05; Fig. 1D) indicating that individual neurons devoid of β-arrestin-2 did not become tolerant to morphine. Analysis of the entire dataset in Figure 1C by 3-way repeated-measures ANOVA revealed a significant main effect of genotype versus overnight morphine treatment versus acute morphine challenge on action potential threshold [F (1, 22) = 5.73; P = 0.03; Table S1], but not on rheobase [F (1,22) = 0.000136; P = 0.90; Table S1], resting membrane potential [F (1,22) = 0.149; P = 0.70; Table S1] or input resistance [F (1,22) = 1.015; P = 0.32; Table S1].

**Figure 1.**
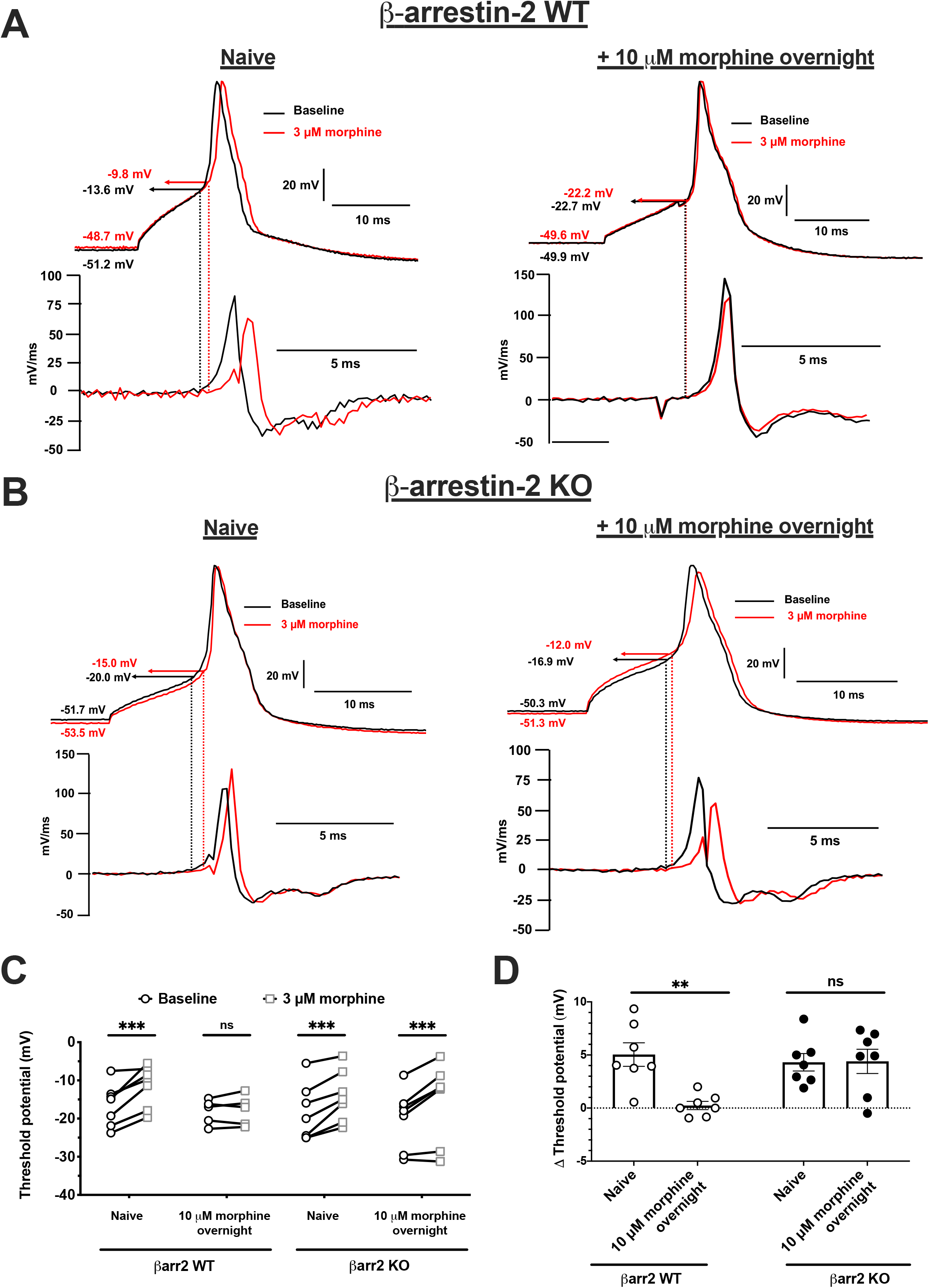
β-arrestin-2 knockout abolishes tolerance to overnight morphine exposure in DRG neurons. Representative short-pulse (10 ms) whole-cell current-clamp traces of naïve and overnight morphine-treated (A) βarr2 WT or (B) βarr2 KO DRG neurons. Action potential is generated at baseline (black) and after 16 minutes of acute 3 μM morphine challenge (red). Threshold potential, which is a measure of neuronal excitability, is extrapolated from the point on the action potential derivative trace, where the action potential differential (dV/dt) > 0. (C) Threshold potentials of βarr2 WT and βarr2 KO DRG neurons at baseline and after bath exposure to 3 μM morphine for up to 16 minutes. Data analyzed by 3-way repeated-measures ANOVA with Bonferroni’s post-test. Significant effect of genotype x overnight morphine treatment x acute morphine challenge on threshold potential [F (1, 22) = 5.73; P = 0.03] was detected. ***P<0.001 vs. baseline and ‘ns’ is not significant (P>0.05). Scatter represents individual cells (D) Threshold potential change from baseline following acute 3 μM morphine challenge for up to 16 minutes. Data analyzed by 2-way ANOVA with Bonferroni’s post-test. Significant main effect of genotype x overnight morphine treatment on threshold potential [F (1, 24) = 7.14; P = 0.01] was detected. Data are mean ± S.E.M. Scatter represents individual cells. **P<0.01 and ‘ns’ is not significant (P>0.05). Naïve βarr2 WT: N = 4 mice n = 7 cells; Overnight-treated βarr2 WT: N = 3, n= 5; Naïve βarr2 KO: N = 5, n = 7; and Overnight-treated βarr2 KO: N = 3, n = 7.

Consequently, these data indicate that short-term morphine tolerance in DRG neurons is mediated by β-arrestin-2.

### 3.2. TRV130 does not produce short-term tolerance in DRG neurons

We next studied the effects of TRV130, a G-protein biased agonist that does not promote the recruitment of β-arrestin-2 on the MOR (DeWire et al., 2013; Pedersen et al., 2020). Neurons isolated from male β-arrestin-2 WT mice were incubated overnight with 200 nM TRV130 and cellular tolerance was evaluated by challenging DRG neurons (membrane capacitance < 30 pF) with 50 nM TRV130 (Fig. 2). In β-arrestin-2 WT neurons incubated in untreated media, acute exposure to 50 nM TRV130 led to statistically significant shifts in threshold potentials to more positive values compared to baseline recordings (−14.0 ± 1.8 mV to −10.4 ± 1.5 mV, P < 0.01; Figs 2A and 2C; Table S2). Following overnight incubation in 200 nM TRV130, neurons challenged with 50 nM TRV130 on Day 2 continued to show statistically significant positive shifts in threshold potential compared to baseline (−12.8 ± 0.9 mV to −10.0 ± 1.0 mV, P < 0.05, Figs. 2B and 2C, Table S2), indicating that overnight TRV130 exposure did not produce tolerance in individual DRG neurons. Analysis of the entire TRV130 dataset in Figure 2C by 2-way repeated-measures ANOVA did not reveal a significant main effect of overnight TRV130 treatment versus acute TRV130 challenge on action potential threshold [F (1,16) = 0.27; P = 0.61; Table S2]. Furthermore, in both naïve and overnight TRV130-treated neurons the mean threshold potential change following acute TRV130 challenge is not significantly different (Naïve: 3.6 ± 1.0 vs. Overnight-treated: 2.8 ± 1.2; Fig. 2D).

**Figure 2.**
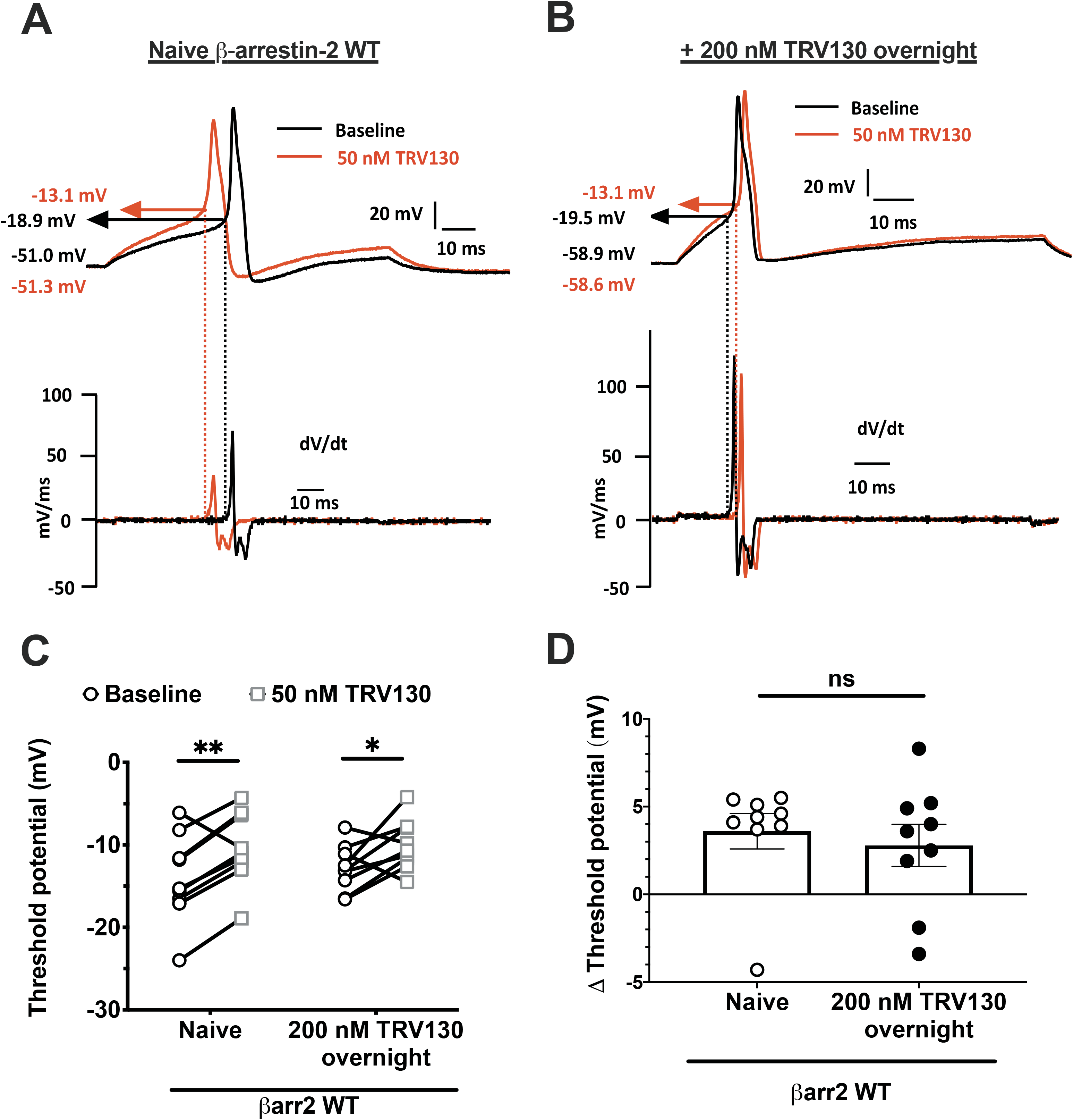
Overnight exposure to TRV130 does not produce tolerance in β-arrestin-2 WT DRG neurons. Representative long-pulse (100 ms) current-clamp traces of (A) naïve and (B) overnight TRV130-treated βarr2 WT DRG neurons. Action potential is generated at baseline (black) and after acute 50 nM TRV130 challenge (red). Threshold potential is extrapolated from the point on the action potential derivative trace, where the action potential differential (dV/dt) > 0. (C) Threshold potentials of naïve and overnight 200 nM TRV130-treated DRG neurons isolated from male βarr2 WT mice at baseline and after acute challenge with 50 nM TRV130. 2-way repeated-measures ANOVA did not detect a main effect of overnight treatment x acute TRV130 challenge on threshold potential [F (1,16) = 0.27; P = 0.61]; however, a significant effect of acute TRV130 challenge [F (1,16) = 16.56; P<0.001] on threshold potential was observed. Data, therefore, analyzed by multiple 2-tailed paired t-tests with two-stage step-up method of Benjamini, Krieger and Yekutieli. The False Discovery Rate was set to 5%. *Adjusted P<0.05 and **adjusted P<0.01 vs. baseline. (D) Threshold potential change from baseline after acute 50 nM TRV130 challenge. ‘ns’ is not significant (P>0.05) by 2-tailed unpaired t-test. Naïve: N = 5, n = 9; and Overnight-treated: N = 2, n = 9. Data are mean ± S.E.M. Scatter represents individual cells.

These studies show that β-arrestin-2-induced desensitization of the MOR mediates short-term opioid tolerance in DRG nociceptors.

### 3.3. Acute antinociceptive tolerance in-vivo is dependent on β-arrestin-2

In order to test if the development of acute antinociceptive tolerance in-vivo is dependent on β-arrestin-2, we utilized a short-term tolerance injection schedule previously published by Bohn et al. (Bohn et al., 2002) and tested antinociception using the warm-water tail withdrawal assay. Morphine was assessed in both β-arrestin-2 WT and KO male mice, while the effects of TRV130 were only assessed in male WT mice as TRV130 prevents β-arrestin-2 activation. Mice were considered drug-responsive if the acute challenge dose significantly increased tail-withdrawal latency. Alternately, mice were deemed tolerant if the acute challenge dose did not significantly increase tail-withdrawal latency. Mice that received saline on Day 1 exhibited antinociception to their respective challenge dose of 10 mg/kg morphine s.c. or 3 mg/kg TRV130 s.c. on Day 2 (Fig. 3). As predicted, WT mice that received a high dose of morphine (100 mg/kg, s.c.) on Day 1 showed tolerance development when challenged on Day 2 (38.5 ± 12.6 %MPE), as compared to saline pre-treated WT controls acutely challenged with 10 mg/kg morphine on Day 2 (85.2 ± 10.1 %MPE, P = 0.007; Fig. 3A). In β-arrestin-2 KO mice however, high dose morphine exposure on Day 1 did not lead to tolerance development when they were challenged on Day 2 (91.2 ± 8.8 %MPE vs. saline controls: 89 ± 11.0, P>0.05; Fig. 3A). Furthermore, 3 mg/kg TRV130 was equally effective as morphine in producing an acute antinociceptive effect (100.0 ± 0.0 %MPE; Fig 3B). A high dose of TRV130 (30 mg/kg, s.c.) given on Day 1 did not lead to tolerance development following a challenge injection on Day 2 (84.72 ± 15.3 %MPE) as compared to acute TRV130 wild type controls (100.0 ± 0.0 %MPE, P>0.05; Fig. 3B).

**Figure 3.**
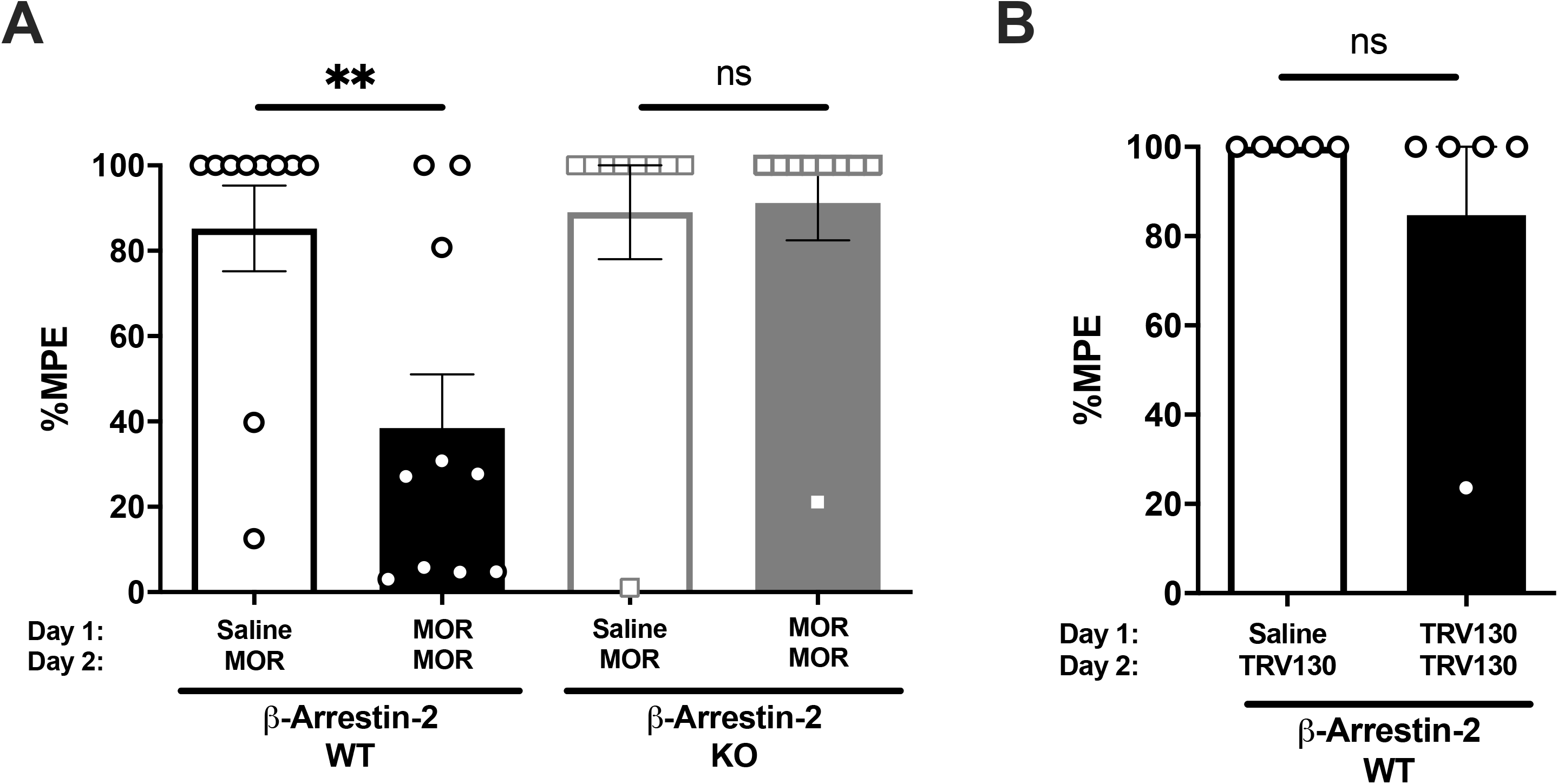
Acute antinociceptive tolerance in mice is mediated by β-arrestin-2. (A) Warm-water tail-withdrawal antinociception in β-arrestin-2 WT or KO mice pre-treated with either saline or 100 mg/kg morphine s.c. on Day 1 and acutely challenged with 10 mg/kg morphine s.c on Day 2. Data are %MPE ± S.E.M represent Day 2 results. Scatter represents individual mice. β-arrestin-2 WT mice: N = 10/group; β-arrestin-2 KO mice: N = 9/group. Data analyzed by 2-way ANOVA with Bonferroni’s post-test. Main effect of Day 1 pre-treatment x genotype on %MPE was significant [F (1, 34) = 5.139; P = 0.03]. ***P<0.01 and ‘ns’ is not significant (P>0.05). (B) Warm-water tail-withdrawal antinociception in β-arrestin-2 WT mice pre-treated with either saline or 30 mg/kg TRV130 s.c. on Day 1 and acutely challenged with 3 mg/kg TRV130 s.c on Day 2. Data are %MPE ± S.E.M and represent Day 2 results. Scatter are individual mice. β-arrestin-2 WT mice: N = 5/group. Data analyzed by 2-tailed unpaired t-test. ‘ns’ is not significant (P>0.05). WT mice injected with 100 mg/kg morphine on Day 1 respond significantly less to acute morphine challenge on Day 2 vs. saline controls, indicating tolerance. Mice devoid of β-arrestin-2 or pre-treated with TRV130 on Day 1 continue to respond to the acute morphine or TRV130 challenge on Day 2, respectively, indicating no acute antinociceptive tolerance development in absence of β-arrestin-2 activation.

Altogether, these results support the previously published finding that β-arrestin-2 mediates acute antinociceptive tolerance at the whole animal level.

### 3.4. Long-term exposure to morphine induces tolerance in DRG neurons independent of β-arrestin-2

We investigated whether tolerance after long-term exposure to morphine is mediated by β-arrestin-2. In order to test this, DRG neurons were collected from male β-arrestin-2 WT or KO mice subcutaneously implanted with a 50 mg morphine pellet for 7 days. As in previous studies, cellular tolerance was assessed by challenging small-diameter DRG neurons (membrane capacitance < 30 pF) with 3 μM morphine in the bath and action potential threshold was used as the indicator of cellular excitability (Fig.4). The 3 μM morphine challenge did not appear to produce a shift in action potential threshold from baseline values of DRG neurons obtained from βarr2 WT mice exposed to 50 mg morphine pellet for 7 days (−22.3 ± 2.3 mV to −22.2 ± 2.2 mV; Figs. 4A and 4C; Table S3). Interestingly, 3 μM morphine also appeared to not alter the threshold potential from baseline of DRG neurons isolated from 7-day 50 mg morphine-pelleted βarr2 KO animals (−17.9 ± 1.8 mV to −19.1 ± 1.7 mV; Figs. 4B and 4C; Table S3). Analysis of the entire dataset by 2-way repeated-measures ANOVA did not detect an effect of acute morphine challenge on threshold potential [F (1, 15) =1.20; P = 0.29]. The ANOVA analysis also did not reveal a significant main effect of genotype versus acute morphine challenge on action potential threshold [F (1, 15) =1.565; P = 0.23]. Furthermore, mean threshold potential change after acute morphine challenge was not significantly different between the WT and KO DRG neurons (P > 0.05 by 2-tailed unpaired t-test; Fig. 4D). No significant effect of genotype and acute morphine challenge on rheobase, resting membrane potential or input resistance was observed (Table S3). Altogether, these data indicated the development of morphine tolerance in DRG neurons from morphine-pelleted βarr2 WT and KO male mice.

**Figure 4.**
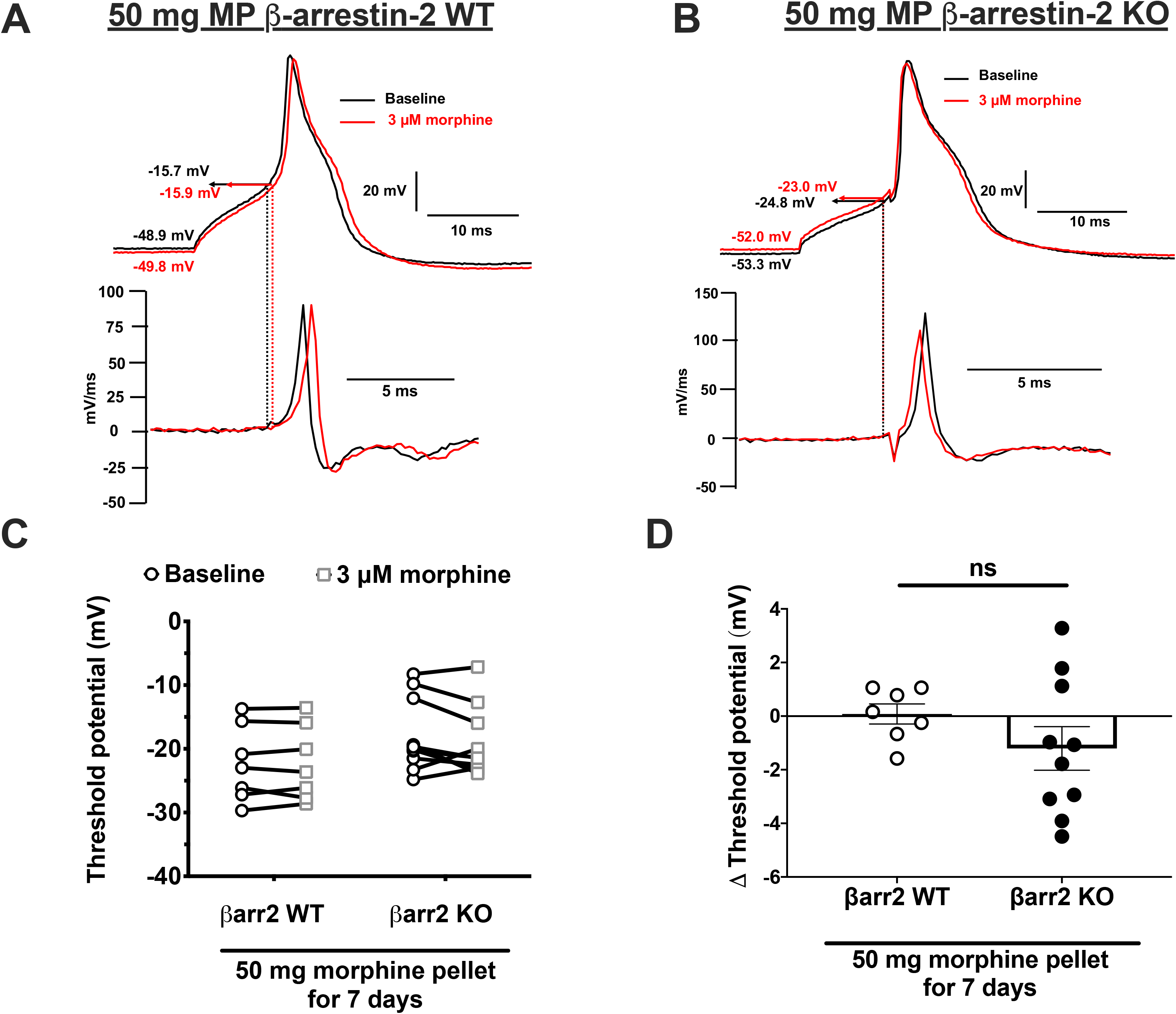
β-arrestin-2 knockout does not prevent tolerance in DRG neurons isolated from chronic morphine-treated mice. Representative current-clamp traces of DRG neurons isolated from (A) βarr2 WT or (B) βarr2 KO male mice treated with 50 mg morphine pellet (MP) for 7 days. Action potential is generated at baseline (black) and after 16 minutes of acute 3 μM morphine challenge (red). Threshold potential is extrapolated from the point on the action potential derivative trace, where the action potential differential (dV/dt) > 0. (C) Threshold potential values of individual neurons from MP βarr2 WT or MP βarr2 KO mice at baseline and after 3 μM morphine challenge for up to 16 minutes. Main effect of genotype x acute morphine challenge on threshold potential was not observed [F (1, 15) =1.57; P = 0.23] by 2-way repeated-measures ANOVA. The test also did not detect an effect of acute morphine challenge on threshold potential [F (1, 15) =1.20; P = 0.29]. (D) Threshold potential change from baseline after acute 3 μM morphine challenge. ‘ns’ is not significant (P>0.05) by 2-tailed unpaired t-test. βarr2 WT: N = 4 mice, n = 7 cells; and βarr2 KO: N = 5, n = 10. Data are mean ± S.E.M. Scatter represents individual cells.

Consequently, these findings suggest that unlike short-term tolerance, β-arrestin-2 is not required for long-term tolerance to morphine in DRG neurons.

### 3.5. Long-term morphine tolerance to antinociception is not altered by β-arrestin-2 deletion in either male or female mice

We investigated whether β-arrestin-2 modulates the development of chronic or “long-term” antinociceptive tolerance in mice. To induce tolerance, male and female mice were implanted subcutaneously with 50 mg or 25 mg morphine pellets, respectively, in accordance with a previous report by our laboratory (Muchhala et al., 2020). We determined the development of tolerance over 7 days by an acute challenge of morphine (10 mg/kg morphine s.c). Response to the challenge dose was tested on Days 1, 3, 4 and 7 post pellet implantation. Pre-injection baseline latency was compared with post-injection latency on these days. Mice sensitive to morphine-induced antinociception exhibited increased tail-withdrawal latencies that were at or close to the 10-second maximum cutoff. Mice were regarded as tolerant if the acute morphine challenge failed to induce antinociception i.e. no statistically significant change (P>0.05) in tail-withdrawal latency from baseline. As predicted, 10 mg/kg morphine increased tail-withdrawal latency of most placebo-pelleted mice (46/49 mice) to the maximum cutoff of 10 seconds (Fig. 5 for males and Fig. 6 for females). In 1-day morphine-pelleted male βarr2 WT and KO mice, the baseline tail-withdrawal latency was at the 10-second maximum cutoff, indicating morphine-induced antinociception (Fig. 5A). The baseline reduced in 3 out of 7 mice in the WT cohort by day 3 indicative of a slow development of tolerance. All βarr2 KO male mice continued to exhibit maximal baseline tail-withdrawal latencies three days after morphine pellet exposure (Fig. 5B). Interestingly, after four days of morphine pellet exposure the antinociceptive effect of the morphine challenge extinguished in both male βarr2 WT and KO animals (P>0.05 vs. baseline by 2-way repeated-measures ANOVA with Bonferroni’s post-test), indicating the development of morphine tolerance despite β-arrestin-2 deletion (Fig. 5C). Tolerance to 10 mg/kg morphine was observed in all male mice (7/7 WT mice and 8/8 KO mice; P>0.05 vs. baseline by 2-way repeated-measures ANOVA with Bonferroni’s post-test) after 7 days of morphine exposure (Fig. 5D). Thus, β-arrestin-2 knockout neither shifted the time-course of morphine tolerance nor prevented the manifestation of morphine tolerance in male mice.

**Figure 5.**
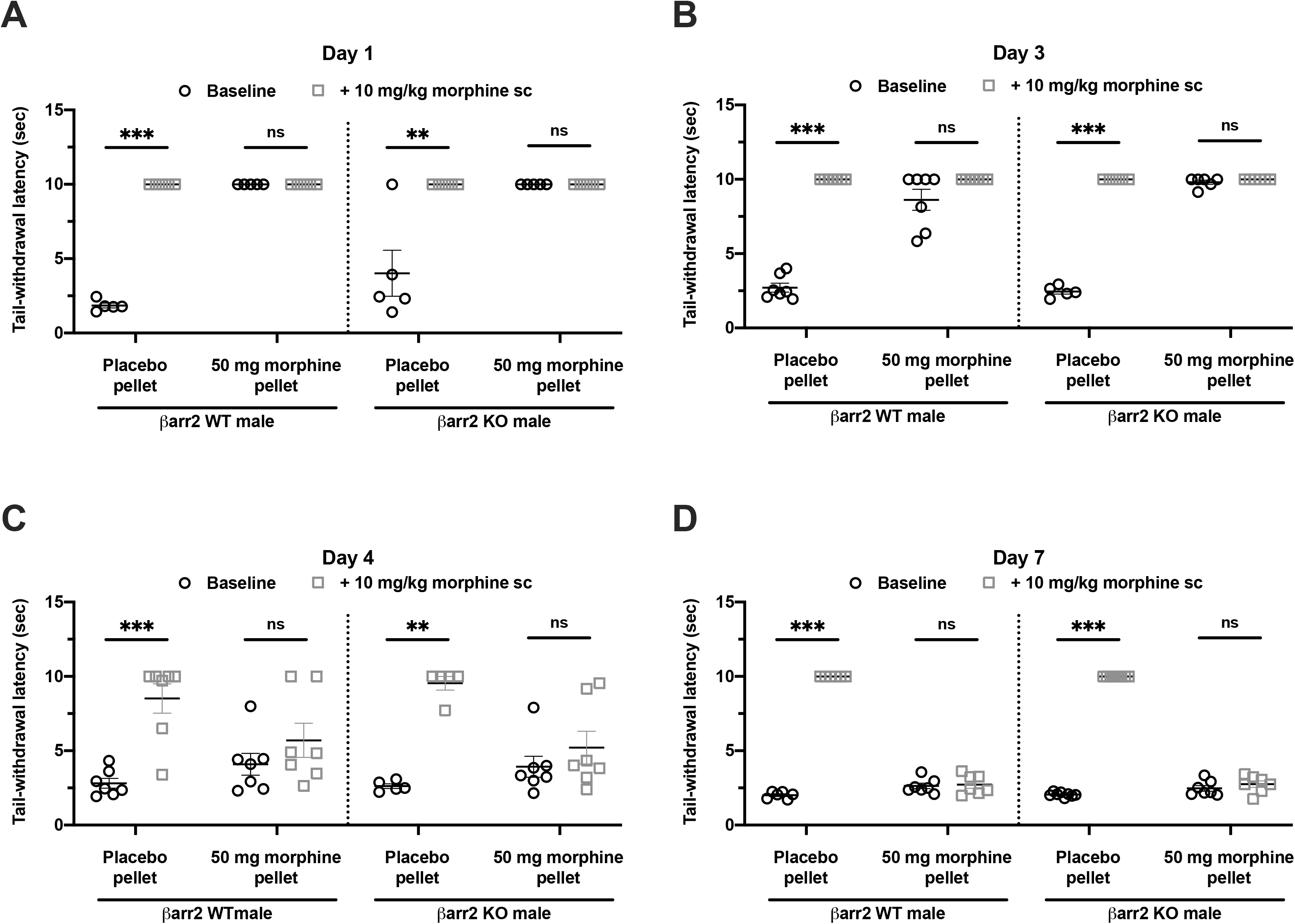
Long-term morphine tolerance develops to tail-withdrawal antinociception in male mice in the absence of β-arrestin-2. Long-term morphine tolerance was evaluated by acutely challenging male βarr2 WT (left) or KO (right) mice implanted with placebo or 50 mg morphine pellet with 10 mg/kg morphine s.c. on days (A) 1, (B) 3, (C) 4 or (D) 7, and tail-withdrawal latencies before and after challenge were compared. Data are mean ± S.E.M. Scatter represents individual animals. **P<0.01, ***P<0.001 and ‘ns’ (not significant, P>0.05) vs. baseline by 2-way repeated-measures ANOVA with Bonferroni’s post-test. Day 1: N=5/group; Day 3: N = 7 (PP WT), 5 (PP KO), 7 (MP WT) and 6 (MP KO); Day 4: N = 7 (PP WT), 5 (PP KO), 7 (MP WT) and 7 (MP KO); Day 7: N = 6 (PP WT), 8 (PP KO), 7 (MP WT) and 7 (MP KO). Both, βarr2 WT and KO mice develop antinociceptive tolerance over 7 days of morphine exposure, indicating that long-term antinociceptive tolerance in male mice is independent of β-arrestin-2.

**Figure 6.**
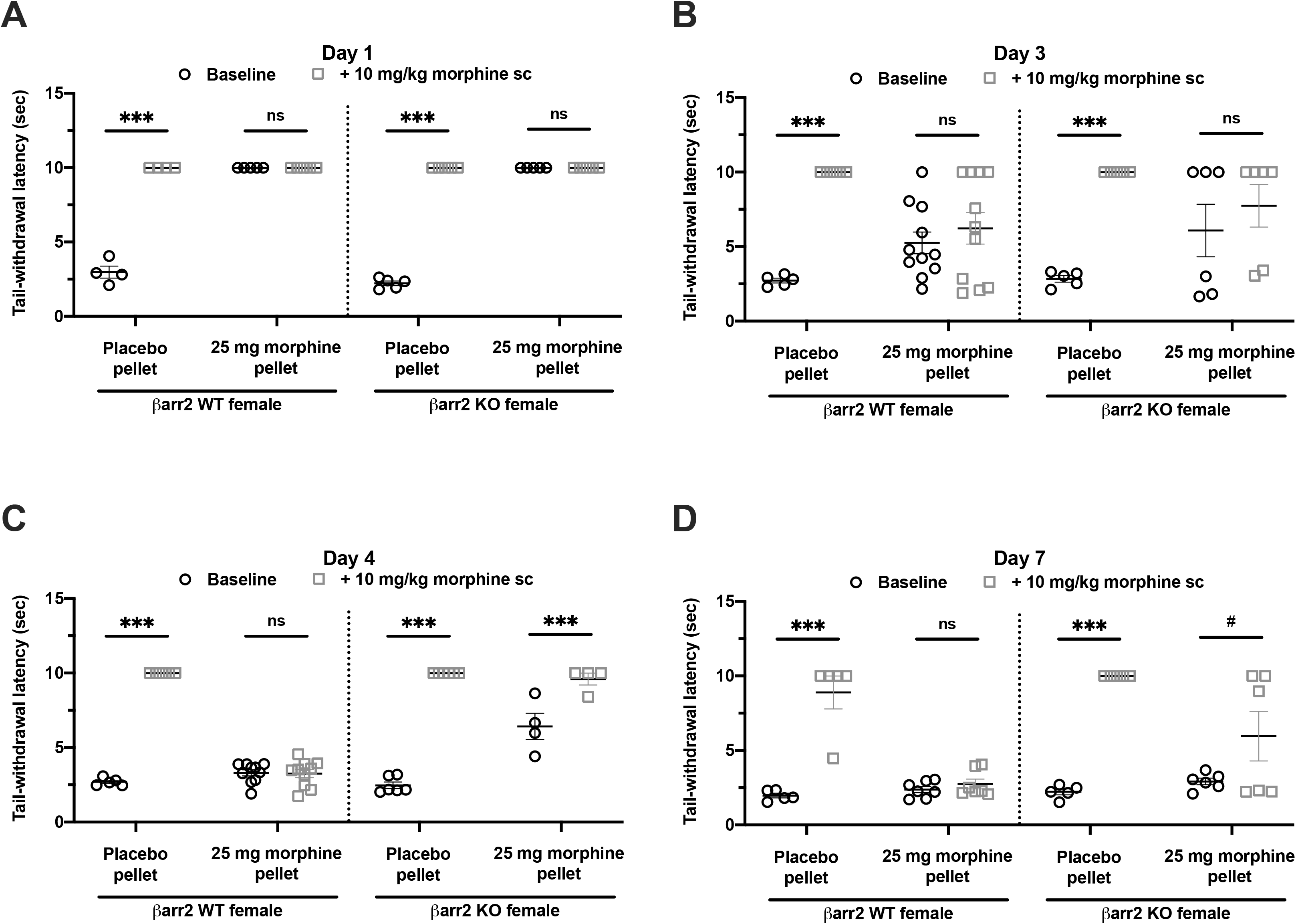
Long-term morphine tolerance develops to tail-withdrawal antinociception in female mice in the absence of β-arrestin-2. Morphine tolerance was evaluated by acutely challenging female βarr2 WT (left) or KO (right) mice implanted with placebo or 25 mg morphine pellet with 10 mg/kg morphine s.c. on days (A) 1, (B) 3, (C) 4 or (D) 7, and tail-withdrawal latencies before and after challenge were compared. Data are mean ± S.E.M. Scatter represents individual animals. ***P<0.001, #P = 0.06 and ‘ns’ (not significant, P>0.05) vs. baseline by 2-way repeated-measures ANOVA with Bonferroni’s post-test. Day 1: N = 4 (PP WT) and rest 5/group; Day 3: N = 5 (PP WT), 5 (PP KO), 11 (MP WT) and 6 (MP KO); Day 4: N = 5 (PP WT), 6 (PP KO), 10 (MP WT) and 4 (MP KO); Day 7: N = 5 (PP WT), 5 (PP KO), 7 (MP WT) and 6 (MP KO). Antinociceptive tolerance develops in both βarr2 WT and KO female mice, indicating that antinociceptive tolerance in female mice is independent of β-arrestin-2.

Similar to observations in male mice, 1-day morphine-pelleted female mice exhibited the maximum 10-second cutoff baseline latency (Fig. 6A). After three days of morphine exposure, 10/11 βarr2 WT and 3/6 KO mice responded in the tail-withdrawal test at baseline, i.e. submaximal baseline tail-withdrawal latencies were observed (Fig. 6B). Acute 10 mg/kg morphine challenge, however, did not significantly increase antinociception compared to baseline in either genotype (Fig. 6B; P>0.05 vs. baseline by 2-way repeated-measures ANOVA). The lack of response to the morphine challenge was also evident in 4-day and 7-day morphine-pelleted female WT mice, indicating the development of morphine tolerance (Figs. 6C and 6D). In contrast, 10 mg/kg morphine produced antinociception in female βarr2 KO mice exposed to morphine for four days (Fig. 6C; P<0.001 vs. baseline by 2-way repeated-measures ANOVA with Bonferroni’s post-test). This trend was also observed in 3/6 7-day morphine-pelleted female βarr2 KO mice. The remaining mice exhibited tolerance to the morphine challenge (Fig. 6D; P =0.06 vs. baseline by 2-way repeated-measures ANOVA). Altogether, these data suggest that β-arrestin-2 might not mediate the development of morphine tolerance in female mice.

Finally, we evaluated whether there were sex differences in antinociceptive tolerance in βarr2 KO mice. A cumulative dose-response to morphine was conducted in male and female mice (Fig. 7). Morphine-induced antinociception was quantitated as % MPE, where 100% MPE represented maximal antinociception. ED_50_ values of 7-day placebo-pelleted male and female KO mice were 2.90 (1.89-4.33) mg/kg and 1.56 (1.25-1.98) mg/kg, respectively. Morphine pre-treatment for 7 days produced a significant rightward shift in the dose response curve of both male and female mice, irrespective of genotype (Fig. 7). The ED_50_ values for 7-day morphine-pelleted male and female mice were 63.9 (42.4-113.7) mg/kg and 41.82 (31.66-55.61) mg/kg, respectively. Decreased potency suggested that morphine tolerance was induced in βarr2 KO mice, irrespective of the sex.

**Figure 7.**
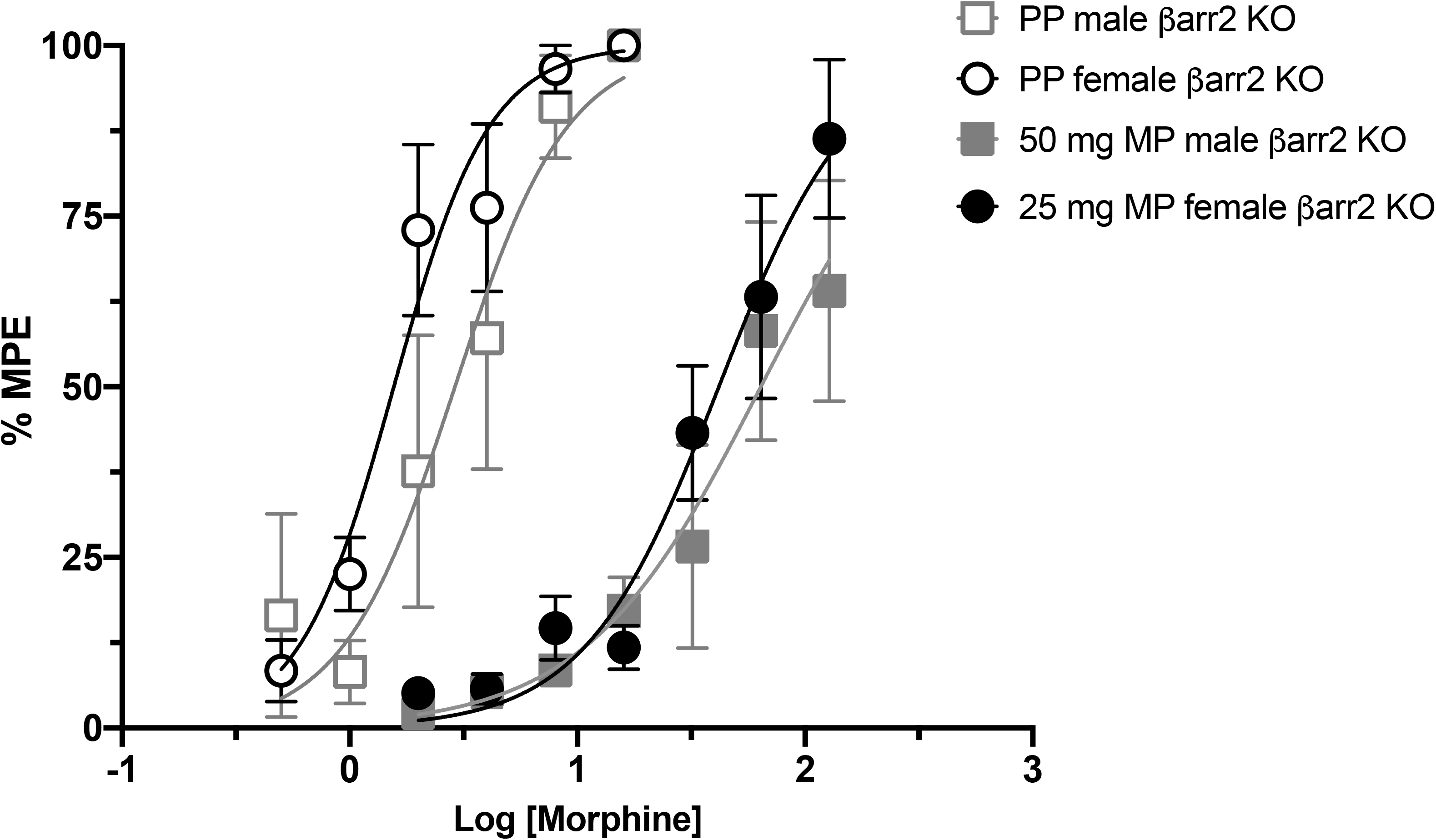
Cumulative morphine dose response curve of male and female β-arrestin-2 KO mice. Cumulative morphine dose response curves of 7-day placebo-pelleted (PP) and morphine-pelleted (MP) male and female mice were compared to evaluate the effect of β-arrestin-2 deletion on warm-water tail-withdrawal antinociception. Each point is %MPE ± S.E.M. Males: N = 6/group. Females: 7/group. ED50 of 50 mg MP male mice is significantly right-shifted compared to PP male mice [MP: 63.9 (42.4-113.7) vs. PP: 2.90 (1.89-4.33)]. Similarly, ED50 of 25 mg MP female mice is significantly right-shifted compared to PP female mice [MP: 41.82 (31.66-55.61) vs. PP: 1.56 (1.25-1.98)]. Thus, antinociceptive tolerance develops in both male and female mice despite the absence of β-arrestin-2

Consequently, these data suggest that β-arrestin-2 does not regulate long-term antinociceptive tolerance in either male or female mice.

## 4. Discussion

The CNS is traditionally considered as the primary site of opioid-induced antinociceptive tolerance and, therefore, research has primarily focused on delineating mechanisms underlying opioid tolerance in centrally-localized MORs. However, systemically-administered opioids can induce analgesia by concomitantly activating MORs at multiple sites along the pain pathway, including the primary afferent neurons of the DRG. MORs expressed on DRG primary afferent neurons are critical for regulating the influx of nociceptive stimuli to the CNS. Activation of peripheral MORs can prevent the sensitization of primary afferent neurons to noxious stimuli and attenuate subsequent CNS events underlying the perception of pain. Alternately, desensitization of MORs on DRG neurons as a result of chronic opioid exposure can render this process inactive. Recent studies have in fact demonstrated that MORs expressed by nociceptive neurons of the DRG profoundly contribute to the induction of antinociceptive tolerance in mice (Chen et al., 2007; Corder et al., 2017). It is therefore critical to investigate mechanisms underlying opioid tolerance in DRG neurons.

In the present study, we observed that acute tolerance in single nociceptive DRG neurons and to tail-withdrawal antinociception is prevented by genetic deletion of β-arrestin-2 or by using TRV130, a G-protein-biased agonist at the μ-opioid receptor. In contrast, long-term morphine tolerance in individual nociceptive DRG neurons and to tail-withdrawal antinociception in either male or female mice develops independently of the β-arrestin-2 pathway. Taken together the findings presented here indicate that the different phases of antinociceptive tolerance are regulated via distinct mechanisms— a rapid β-arrestin-2-dependent mechanism that mediates acute tolerance, and a slow β-arrestin-2-independent mechanism that underlies long-term tolerance—in both mice and individual DRG neurons critical in the initiation of nociceptive stimuli.

The development of G-protein-biased MOR agonists that preferentially reduce β-arrestin-2 activation have generated enthusiasm within the field and opened up new avenues for the treatment of pain with reduced risks such as tolerance (Madariaga-Mazón et al., 2017; Siuda et al., 2017). In fact, the G-protein biased MOR agonist, Olinvyk (TRV130), was recently approved by the FDA for short-term intravenous use in hospitals (U.S. Food and Drug Administration, 2020). The findings reported in the present study implicate that antinociceptive tolerance can develop in the absence of β-arrestin-2 activation. Consequently, these findings raise questions about the usefulness of G-protein-biased agonists to mitigate opioid-induced analgesic tolerance.

β-arrestins are universally expressed multi-functional adaptor proteins that prevent the coupling of GPCRs to cognate G-proteins through steric hindrance and consequently, signal transduction through membrane-delimited mechanisms is disrupted (Lohse et al., 1990). Multiple studies have implicated that this β-arrestin-2 pathway is the basis of opioid-induced analgesic tolerance (Bohn et al., 2000; DeWire et al., 2013; Grim et al., 2020; Manglik et al., 2016; Wang et al., 2016; Yang et al., 2011). Consistent with these findings, in the present study we too observed that antinociceptive tolerance in mice and cellular tolerance in nociceptive DRG neurons, specifically acute tolerance (Williams et al., 2013), is mediated by β-arrestin-2. Interestingly, in the current study tolerance induced after long-term morphine exposure (7 days) in both mice and DRG neurons was not contingent on β-arrestin-2. Altogether, the data suggests that β-arrestin-2 mediates only the acute but not the long-term phase of tolerance at the MOR. Indeed, Bohn et al. have previously reported that the onset of antinociceptive tolerance in the warm-water tail-withdrawal assay in mice is delayed but not prevented in the absence of β-arrestin-2 (Bohn et al., 2002). In contrast, functional deletion of β-arrestin-2 in rats prevented the development of chronic morphine tolerance to warm-water tail-withdrawal antinociception (Wang et al., 2016; Yang et al., 2011). This discrepancy could be due to species differences or disparate tolerance development models, for example daily injections vs. continuous-release subcutaneous pellets. It is well-known that the extent of tolerance produced by subcutaneous pellets is significantly greater than other techniques (Dighe et al., 2009) and therefore, tolerance produced by intermittent morphine injections might be more readily reversed compared to tolerance produced by morphine pellets.

Chronic exposure to morphine is known to induce bacterial translocation and inflammation of the gut wall and these processes have been previously implicated in the development of opioid tolerance (Kang et al., 2017; Komla et al., 2019; Meng et al., 2013; Mischel et al., 2018). Antinociceptive tolerance in mice was attenuated by modulating the gut microbiome with antibiotics or probiotics (Kang et al., 2017; Mischel et al., 2018; Zhang et al., 2019). Altering the gut microbiome also prevented morphine tolerance at the single-cell level in DRG neurons (Kang et al., 2017; Mischel et al., 2018). Conversely, colonic inflammation enhanced the rate of morphine tolerance to antinociception (Komla et al., 2019). It is therefore, possible that long-term tolerance to antinociception in mice and in DRG neurons might be mediated by changes induced within the gut microbiome.

Previous studies have reported sex differences in opioid analgesia and tolerance in both humans and rodents (Bodnar and Kest, 2010; Craft et al., 1999; Kalinichev et al., 2001; Kasson and George, 1984; Lee and Ho, 2013; Mousavi et al., 2007). However, it was not known whether sex is an important variable influencing the role of β-arrestin-2 in the mechanism of antinociceptive tolerance. In the present study, we find that antinociceptive tolerance in the warm-water tail-withdrawal assay develops in both male and female mice even in the absence of β-arrestin-2, implicating that sex hormones might not interact with the β-arrestin-2 pathway to mediate antinociceptive tolerance in mice.

It is noteworthy that in the present study relatively high doses of morphine were used to investigate tolerance development *in vivo* in mice (25 mg and 50 mg morphine pellet for 7 days in female and male mice, respectively) and *in vitro* in DRG neurons (3 μM morphine for acute challenge and 10 μM morphine for overnight treatment). Previously, it has been reported in 8-week old male C57Bl/6NCr mice that a 25 mg morphine pellet produces a peak plasma morphine concentration of 2695.3 ± 785.1 ng/mL (~3.5 μM morphine) 24 hours after pellet implantation, which progressively decreased to 229.5 ± 92.5 ng/mL after 7 days (~0.4 μM morphine) (McLane et al., 2017). In humans, therapeutic doses of morphine have been reported to produce serum concentrations of 14.7-70.4 ng/mL (Netriova et al., 2006), whereas in overdose cases, wide-ranging plasma concentrations from of 113 ng/mL to 4660 ng/mL have been detected (Meissner et al., 2002; Ozaita et al., 2002). Thus, while the amount of morphine delivered to mice and isolated neurons in our experiments is on the higher end of what patients might receive in the clinic, it is comparable to what might be observed in opioid abusers.

In conclusion, the findings presented in this study implicate that antinociceptive tolerance involving peripheral MORs in the DRG is mediated by two disparate mechanisms—β-arrestin-2-dependent and-independent pathways—that are engaged during different phases of opioid exposure. This suggests that the contribution of β-arrestin-2-induced desensitization of MORs in the molecular mechanism of tolerance is more intricate and that tolerance might be mediated by other means such as microbial dysbiosis in the gut. Importantly, these findings help inform the clinical utility of chronic exposure to G-protein-biased agonists over conventional opioids for pain management and highlights considerations for the likelihood of developing analgesic tolerance.

## Acknowledgments

The authors wish to thank David Stevens and Dr. Krista Scoggins for their technical assistance. This work was supported by the National Institute of Health grants: P30 DA033934, R01 DA036975, R01 DA024009.

