## Supplemental tables for "Role of β-arrestin-2 in short- and long-term opioid tolerance in the dorsal root ganglia"

|  |  | Threshold potential (mV) | Rheobase (pA) | Resting membrane potential (mV) | Input resistance (MΩ) | Cell capacitance (pF) |
| --- | --- | --- | --- | --- | --- | --- |
| **Naïve βarr2 WT** | Baseline | -16.5 ± 2.1 | 152 ± 19 | -52.3 ± 1.0 | 269 ± 36 | 14.9 ± 1.8 |
|  | + 3 µM morphine | -11.4 ± 2.0*** | 156 ± 22 | -51.4 ± 1.6 | 266 ± 37 |  |
| **Overnight morphine-treated βarr2 WT** | Baseline | -18.2 ± 1.5 | 162 ± 66 | -48.9 ± 3.0 | 315 ± 88 | 20.0 ± 2.0 |
|  | + 3 µM morphine | -17.9 ± 1.8 | 159 ± 53 | -46.6 ± 3.8 | 290 ± 86 |  |
| **Naïve βarr2 KO** | Baseline | -18.4 ± 2.8 | 98 ± 17 | -53.9 ± 3.6 | 413 ± 85 | 16.1 ± 3.2 |
|  | + 3 µM morphine | -14.1 ± 2.5*** | 99 ± 16 | -53.7 ± 4.2 | 451 ± 101 |  |
| **Overnight morphine-treated βarr2 KO** | Baseline | -19.9 ± 3.0 | 109 ± 27 | -52.7 ± 2.1 | 420 ± 96 | 15.9 ± 2.1 |
|  | + 3 µM morphine | -15.5 ± 3.9*** | 103 ± 24 | -52.3 ± 2.2 | 491 ± 121 |  |

**Supplementary Table 1 (Related to Figure 1). Electrophysiological properties of DRG neurons.** Data are mean ± S.E.M. 3-way repeated-measures ANOVA found a significant effect of genotype x overnight morphine treatment x acute morphine challenge on threshold potential [F (1,22) = 5.73; P = 0.03]. ***P<0.001 vs. baseline by Bonferroni’s multiple comparisons test. 3-way repeated-measures ANOVA did not detect a main effect of genotype x overnight morphine treatment x acute morphine challenge on rheobase [F (1,22) = 0.000136; P = 0.90], resting membrane potential [F (1,22) = 0.149; P = 0.70] or input resistance [F (1,22) = 1.015; P = 0.32]. It is noteworthy that all neurons tested have cellular capacitance < 30 pF.

|  |  | Threshold potential (mV) | Rheobase (pA) | Resting membrane potential (mV) | Input resistance (MΩ) | Cell capacitance (pF) |
| --- | --- | --- | --- | --- | --- | --- |
| **Naïve βarr2 WT** | Baseline | -14.0 ± 1.8 | 56 ± 17 | -50.9 ± 2.0 | 1132 ± 237 | 15.0 ± 0.9 |
|  | + 50 nM TRV130 | -10.4 ± 1.5** | 40 ± 12 | -48.0 ± 1.3 | 1155 ± 173 |  |
| **Overnight TRV130-treated βarr2 WT** | Baseline | -12.8 ± 0.9 | 31 ± 7 | -46.7 ± 2.9 | 1629 ± 163 | 14.5 ± 0.7 |
|  | + 50 nM TRV130 | -10.0 ±1.0* | 41 ± 11 | -47.4 ± 2.5 | 1351 ± 288 |  |

**Supplementary Table 2 (Related to Figure 2). Electrophysiological properties of DRG neurons.** Data are mean ± S.E.M. 2-way repeated-measures ANOVA did not detect a main effect of overnight TRV130 pre-treatment x acute TRV130 challenge on threshold potential [F (1, 16) = 0.27; P = 0.61], however, a significant effect of acute TRV130 challenge [F (1,16) = 16.56; P<0.001] on threshold potential was observed. Data were, therefore, analyzed by multiple 2-tailed paired t-tests with two-stage step-up method of Benjamini, Krieger and Yekutieli. The False Discovery Rate was set to 5%. *Adjusted P<0.05 and **adjusted P<0.01 vs. baseline. No significant effect of overnight TRV130 pre-treatment x acute TRV130 challenge on rheobase [F (1,16) = 4.046; P = 0.06], resting membrane potential [F (1,16) = 4.109; P = 0.06] or input resistance [F (1,16) = 0.593; P = 0.45] was observed by 2-way repeated-measures ANOVA. It is noteworthy that all neurons tested have cellular capacitance < 30 pF.

|  |  | Threshold potential (mV) | Rheobase (pA) | Resting membrane potential (mV) | Input resistance (MΩ) | Cell capacitance (pF) |
| --- | --- | --- | --- | --- | --- | --- |
| **7-day MP βarr2 WT** | Baseline | -22.3 ± 2.3 | 146 ± 44 | -56.9 ± 1.6 | 339 ± 66 | 13.0 ± 1.8 |
|  | + 3 µM morphine | -22.2 ± 2.2 | 161 ± 67 | -58.2 ± 1.7 | 342 ± 58 |  |
| **7-day MP βarr2 KO** | Baseline | -17.9 ± 1.8 | 113 ± 17 | -54.8 ± 2.1 | 359 ± 38 | 13.3 ± 1.0 |
|  | + 3 µM morphine | -19.1 ± 1.7 | 111 ± 15 | -55.9 ± 2.3 | 380 ± 46 |  |

**Supplementary Table 3 (Related to Figure 3). Electrophysiological properties of DRG neurons.** Data are mean ± S.E.M. 2-way repeated-measures ANOVA did not detect a main effect of genotype x acute morphine challenge on threshold potential [F (1, 15) =1.565; P = 0.23], rheobase [F (1,15) = 0.698; P = 0.42], resting membrane potential [F (1,15) = 0.023; P = 0.88] or input resistance [F (1,15) = 0.271; P = 0.61]. It is noteworthy that all neurons tested have cellular capacitance < 30 pF.
